# Converting networks to predictive logic models from perturbation signalling data with CellNOpt

**DOI:** 10.1101/2020.03.04.976852

**Authors:** Enio Gjerga, Panuwat Trairatphisan, Attila Gabor, Hermann Koch, Celine Chevalier, Francesco Ceccarelli, Aurelien Dugourd, Alexander Mitsos, Julio Saez-Rodriguez

## Abstract

**Motivation:** The molecular changes induced by perturbations such as drugs and ligands are highly informative of the intracellular wiring. Our capacity to generate large data-sets is increasing steadily as new experimental approaches are developed. A useful way to extract mechanistic insight from the data is by integrating them with a prior knowledge network of signalling to obtain dynamic models. Logic models scale better with network size than alternative kinetic models, while keeping the interpretation of the model simple, making them particularly suitable for large datasets.

**Results:** CellNOpt is a collection of Bioconductor R packages for building logic models from perturbation data and prior knowledge of signalling networks. We have recently developed new components and refined the existing ones. These updates include (i) an Integer Linear Programming (ILP) formulation which guarantees efficient optimisation for Boolean models, (ii) a probabilistic logic implementation for semi-quantitative datasets and (iii) the integration of MaBoSS, a stochastic Boolean simulator. Furthermore, we introduce Dynamic-Feeder, a tool to identify missing links not present in the prior knowledge. We have also implemented systematic *post-hoc* analyses to highlight the key components and parameters of our models. Finally, we provide an R-Shiny tool to run CellNOpt interactively.

**Availability:** R-package(s): https://github.com/saezlab/cellnopt

**Supplementary information:** Supplemental Text.

## 1. Introduction

Logic networks are one of the easiest to understand modelling frameworks. They capture the mechanistic relationship between molecular entities by logic gates (Le Novère 2015). Due to their simplicity, they are highly scalable and widely applied. We have previously introduced CellNOpt, a framework to build predictive logic models of signalling pathways by training a prior knowledge network (PKN) to biochemical data obtained from perturbation experiments (Terfve et al. 2012). CellNOpt features different formalisms ranging from Boolean logic to logic-based ordinary differential equations (logic-ODE). These different formalisms have particular strengths and weaknesses, with a general trade-off between detail and coverage (Terfve et al. 2012). The value of CellNOpt has been recently demonstrated on several applications (Table S1).

Over the past years, we have continuously added new features and upgrades to the packages to enhance the functionality and computational efficiency of CellNOpt (Fig. 1). The current toolkit covers a unique set of features (see Table S2). Furthermore, for alternative analyses, CellNOpt supports importing from and exporting to other tools via the Cytoscape’s SIF and the SBMLQual (Chaouiya et al. 2013) formats.

**Fig.1.:**
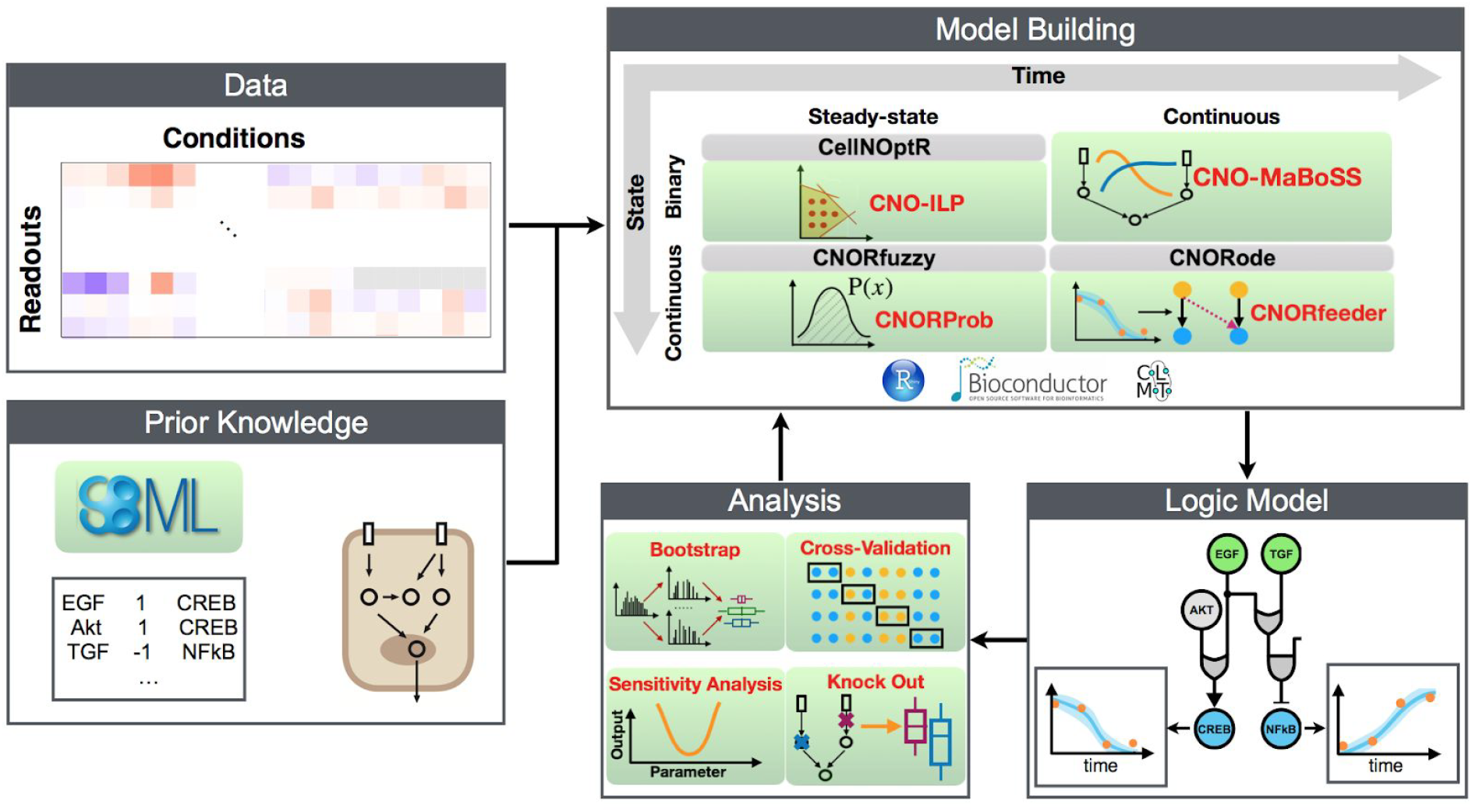
CellNOptR pipeline with packages and features (new implementations highlighted in green). Perturbation data is combined prior knowledge of signalling and CellNOptR is used to contextualise the regulatory signalling interactions.

## 2. Summary of new features

### 2.1. CellNOpt-ILP

For the training of Boolean logic networks, we developed an integer linear programming (ILP) formulation based on (Mitsos et al. 2009). This allows us to optimise large-scale networks orders of magnitude faster than the built-in genetic algorithm. Furthermore, CellNOpt-ILP utilises a CPLEX solver that guarantees global optimality if/once it reaches the solution. The method can enumerate equivalently good solutions based on the objective function values, which helps identify uncertainty in the inferred network structure. We illustrate the functionality of CellNOpt-ILP on several case studies (Text S1).

### 2.2. CellNOpt-MaBoSS

MaBoSS (Stoll et al. 2012) performs asynchronous stochastic simulations of logic Boolean models using the Gillepsie algorithm. It was integrated with CellNOpt to optimise Boolean networks where nodes are represented by their state probability with respect to time (Text S2).

### 2.3. CNORprob

To optimise logic networks with semi-quantitative states (between 0 and 1) at quasi-steady-state, we offer the new CNORprob package. CNORprob is an R-implementation of the Matlab-based toolbox FALCON for probabilistic Boolean logic (Landtsheer et al., 2017). Compared to the CNORfuzzy packages which is based on fuzzy logic (Terfve et al. 2012), CNORprob offers a simpler interpretation as this framework contains only one parameter per edge that has an equivalent probabilistic meaning (Text S3).

### 2.4. Dynamic-Feeder

Our knowledge of pathways is incomplete. Dynamic-Feeder is a method to identify missing links in the PKN and provides candidates to fill such gaps (Fig. S3). It combines data-driven network inference with a protein-protein interaction network to find missing elements. For the latter, we use primarily OmniPath (Türei et al. 2016), an integrated resource of curated pathways’ databases. Dynamic-Feeder generalizes our previous tool CNORfeeder (Eduati et al. 2012) to time-course data with a logic ordinary differential equations (ODE) formalism (Text S4).

### 2.5. Post Hoc analysis

After optimised logic-models are obtained, users can analyse them systematically. These analyses include bootstrapping, cross-validation, parameter sensitivity analysis, and estimation of node and edge essentiality by removing them (knockout). This allows us to observe how sensitive specific proteins/interactions to perturbations are (Text S5).

### 2.6. Shiny application

To use CellNOpt without coding, we offer an interactive version of the packages as an R-Shiny application (Text S6).

## 3. Acknowledgements

The work was supported by the European Union’s H2020 program (675585 Marie-Curie ITN ‘‘SymBioSys’’); JRC for Computational Biomedicine, which is partially funded by Bayer; and the Innovative Medicines Initiative 2 Joint Undertaking under grant agreement no. 116030 (TransQST). We thank Christian Holland for his assistance on setting up the ShinyCNOR tool online and Nicolas Palacio Escat for his help in designing Fig. 1.

