## Supplemental Text for "Converting networks to predictive logic models from perturbation signalling data with CellNOpt"

**Supplementary Figures:**

Figure S1 Genetic Algorithm vs Integer Linear Programming 8
Figure S2 Answer Set Programming vs Integer Linear Programming 8
Figure S3 Dynamic-Feeder Pipeline 13
Figure S4 ShinyCNOR - Loading inputs 18
Figure S5 ShinyCNOR - Network preprocessing 19
Figure S6 ShinyCNOR - CNORprob analysis 20
Figure S7 ShinyCNOR - CNORode analysis 21

**Supplementary Tables**

Table S1 Recent CellNOpt applications 1
Table S2 Features of different logic modelling methods 2
Table S3 Comparison between ILP, ASP and GA for different models 7
Table S4.1 Performance of CellNOpt-MaBoSS to tested software (Case study 1) 10
Table S4.2 Performance of CellNOpt-MaBoSS to CNORode (Case study 2) 11

**Supplementary Texts:**

Text S1 CellNOpt-ILP 4
Text S2 CellNOpt-MaBoSS 9
Text S3 CNORprob 11
Text S4 Dynamic-Feeder 12
Text S5 Post-Hoc Analysis 16
Text S6 ShinyCNOR 17
**References 21**

### Supplementary tables:

**Table S1:** Recent studies where CellNOpt was used. For more applications, please visit http://www.cellnopt.org/.

| Recent CellNOpt applications | |
|---|---|
| (Eduati et al. 2020) | Application to micro-fluidics based perturbation data from biopsies to build logic-ODE models of signalling pathways for individual patients. CellNOpt was used to uncover differences in specific regulatory mechanisms of the apoptosis pathways and to predict patient-specific promising treatments that were validated. |
| (Tognetti et al. 2020, BioRxiv) | Application to mass cytometry single-cell data from breast cancer cell-lines upon stimulation by EGF in the presence of a panel of kinase inhibitors. CellNOpt was used to construct logic-based models of cell signalling in these cell lines. The generated mechanistic signalling network models were predictive of resistance and sensitivity of cell lines to various drugs. |
| (Eduati et al. 2017) | Application to antibody-based phospho-proteomics data obtained from various colorectal cancer cell lines. CellNOpt was used to build cell line-specific dynamic logic-ODE models of underlying signalling networks. Model parameters representing pathway dynamics were used as features to predict drug sensitivities. |
| (Traynard et al. 2017) | Application to a signalling network in prostate cancer. CellNOpt was used to train this network over published phospho-proteomics datasets describing the response of prostate cancer cell lines to a panel of ligands and protein inhibitors. This application highlighted possible molecular mechanisms that drive the androgen-independent growth and androgen-mediated signalling in prostate cancer. |

**Table S2:** Many computational methods have been developed for the logic modelling of biological networks. They can provide qualitative (*GINsim* - Chaouiya, Naldi, and Thieffry 2012, Naldi et al. 2018; *BoolNet* - Müssel, Hopfensitz, and Kestler 2010; ViSiBool - Schwab et al. 2018; *Genetic Network Analyzer* - Batt et al. 2012; *OptimusQual* - Dorier et al. 2016; *PRUNET* - Rodriguez et al. 2015) and quantitative (Stoll et al. 2017; *FALCON* - Landtsheer et al. 2017; *SQUAD* - Di Cara et al. 2007; *optPBN* - Trairatphisan et al. 2014; *BooleanNet* - Albert et al. 2008; *Odefy* - Krumsiek et al. 2010; *Cell Collective* - Helikar et al. 2012; *BioModel Analyzer* - Benque et al. 2012; *CellNOpt* - Terfve et al. 2012) description of a signalling network.

| | ***Simulation w/ continuous states*** | | ***Simulation w/ Boolean states*** | | | | |
|---|---|---|---|---|---|---|---|
| **Tool** | ***Continuous in time*** | ***Discrete in time*** | ***Synchronous updates*** | ***Asynchronous updates*** | ***Graphical User Interface*** | ***Import/export with standards (SBMLqual)*** | ***Model fitting / parameter estimation*** |
| **CellNOpt** (Terfve et al. 2012) | [green] | [green] | [green] | [green] | [green] | [green] | [green] |
| ***GINsim*** (Chaouiya, Naldi, and Thieffry 2012) | | [green] | [green] | [green] | [green] | [green] | |
| ***MaBoSS*** (Stoll et al. 2012) | [green] | [green] | | [green] | | [green] | |
| ***FALCON*** **(**Landtsheer et al. 2017**)** | | [green] | [green] | | [green] | | [green] |
| ***BoolNet*** (Müssel, Hopfensitz, and Kestler 2010) | | [green] | [green] | [green] | | [green] | |
| ***BooleanNet*** (Albert et al. 2008) | [green] | [green] | [green] | [green] | | | |
| **SQUAD** **(**Di Cara et al. 2007**)** | [green] | | | | [green] | | |
| ***optPBN*** **(**Trairatphisan et al. 2014**)** | | [green] | [green] | | | | [green] |

| |
|---|
| ***OptimusQual*** (Dorier et al. 2016) |
| ***ViSiBool*** (Schwab et al. 2018) |
| ***GNA*** (Batt et al. 2012) |
| ***PRUNET*** (Rodriguez et al. 2015) |
| ***Odefy*** (Krumsiek et al. 2010) |
| ***Cell Collective*** (Helikar et al. 2012) |
| ***BMA*** (Benque et al. 2012) |

### Supplementary Text:

### Text S1. CellNOpt-ILP

#### S1.1. Introduction

The problem of training Boolean network models was originally formulated as a non-linear optimisation problem. This task can be solved with stochastic search methods such as a Genetic Algorithm (GA) (Saez-Rodriguez et al. 2009). The solution is often non-unique, i.e. models with different structures can fit the data equally well. The GA finds the models that fit the data with the minimal number of reaction terms. However, biological systems are often not sparse and modellers are also interested in a set of alternative solutions, or when possible, enumerating all the possible solutions that are in agreement with the data. Although GA finds the optimum of objective functions, there is no guarantee that this local optimum is also a global optimum.

Declarative problem-solving technique using Answer Set Programming (ASP) via Cell ASP Optimiser (CASPO) method has been developed for Boolean networks to provide the complete set of optimal models (Guziolowski et al. 2013).

An alternative approach is to solve the optimisation problem using an Integer Linear Programming (ILP) formulation. Like CASPO, the ILP implementation of CellNOpt (CellNOpt-ILP) can provide multiple feasible solutions and it can guarantee that the set of solutions is near optimal under the given tolerance if/when the optimality has been reached. The performances of GA, ASP and ILP have been investigated and compared with respect to the optimisation performance, the predictive capability of the optimised models, and computational effort (see S1.3.2 below).

#### S1.2. Implementation

We have implemented in R language the Integer Linear Programming (ILP) formulation of Boolean model training as presented in (Mitsos et al. 2009) within the CellNOptR framework. The ILP formulation consists of an objective function aimed at minimizing the weighted error between simulations and data. In addition to that, a set of linear constraints (Mitsos et al. 2009) determines the set of feasible interactions present in the model. Based on this formulation, we have implemented new functionalities within the CellNOptR R package to retrieve multiple alternative top-scoring solutions within a tolerance. The ILP problems were solved via the IBM ILOG CPLEX Optimization Studio. The diversity of these solutions can also be controlled through CPLEX settings.

### S1.3. Differences between ILP, GA and ASP

#### S1.3.1. Objective Function

There are differences in the formulation of the objective function between the ILP and the GA/ASP. In both cases, the objective function consists of two parts: an error term $\theta^f$ and a size penalty term $\theta^s$ which ensures that simpler models are preferred over more complicated ones if they explain the data equally well.

Let $x_j^k$ and $x_j^{k,m}$ be the predictions and measurements for each species $j$ in experimental condition $k$. Let also $n_s$ and $n_e$ be the number of species in the PKN and the number of experiments.

In GA and ASP the error term of the objective function is defined through a Mean Squared Error (MSE) function:

$$\theta^f_{GA,ASP} = \frac{1}{n_e * n_s} \sum_{k=1}^{n_e} \sum_{j=1}^{n_s} (x_j^k - x_j^{k,m})^2 , \qquad (1)$$

In contrast, in the ILP formulation the objective function is the sum of absolute differences:

$$\theta^f_{ILP} = \sum_{k=1}^{n_e} \sum_{j=1}^{n_s} | x_j^k - x_j^{k,m} | \qquad (2)$$

In (Mitsos et al. 2009) it has been shown that while the objective functions for GA and ILP are not the same, they still yield the same optimum. In both cases, the optimisation problem is a minimisation problem and we have only shown the part of the objective function that minimises the mismatch between model simulations and observations. Both objective functions have an additional term that penalises the size of the network $\theta^s = \sum_{i=1}^{n_r} \lambda y_i$, where $y_i \in \{0, 1\}$ represents the presence of interaction $i$ and $n_r$ is the total number of interactions in the PKN. $\lambda$ is a size penalty factor.

#### S1.3.2. Efficiency

We compared the optimisation efficiencies of ILP against the Genetic Algorithm/Answer Set Programming (GA/ASP) in terms of computational time and their ability to reach an optimal solution (Fig. S1, Fig. S2). We tested the algorithms on 7 case-studies with increasing complexity: The *ToyPCB* (Morris et al. 2011) and the *ToyMSB2009* (Saez-Rodriguez et al. 2009) models of TGFalpha and TNFalpha signaling pathways; The *ToyPB* (Chaouiya et al. 2013) models the interaction between the MAPK and NFkB cascades; The *LiverDREAM* (Saez-Rodriguez et al. 2009) case-study was part of the DREAM4 - In Silico Network challenge (Chun et al. 2011). *ExtLiverBMC2012* (Terfve et al. 2012) which uses phosphorylation data obtained from a human hepatocellular cell-line (the HepG2) after perturbation with inhibitory drugs; *ExtLiverPCB* (Morris et al. 2011) which is a larger version of *ExtLiverBMC2012*; and *ExtLiverPriHu-MCP2010* (Alexopoulos et al. 2010) case in which phosphorylation levels of several intra-cellular proteins were measured after exposing HepG2 cells to various growth factors, cytokines and small molecule kinase inhibitors. More details about each case-study can be found in Table S3.

For case studies in which all algorithms found identical optimal models, we compared the computation time needed for the solution of the problem. Figure S1 and Figure S2 show the computational times of the different algorithms for models as a function of the number of edges in the PKN. Several observations can be made (Table S3): Firstly, the computation time increases with the number of edges in the preprocessed PKN. Secondly, the computation time for CellNOpt-ILP is orders of magnitude lower than for the Genetic algorithm for all the cases when we retrieve only one solution. More than that, the ILP is able to retrieve the solutions with the best possible optimisation score. Finally, if we compare ILP with CASPO, we observe that CASPO outperforms ILP for the smaller models (*LiverDREAM*, *ToyPB*, *ToyPCB*). However, these differences are fractions of a second. In contrast, for larger models (*ExtLiverPriHu-MCP2010*, *ExtLiverPCB* and *ExtLiverBMC2012*), ILP is more efficient than CASPO. In particular, for the *ExtLiverPriHu-MCP2010* case, CASPO could not retrieve the solutions after 72 hours, after which the optimisation was interrupted. The number of solutions asked to be retrieved by the ILP when compared to CASPO was the most diverse 100 out of the 1000 computed by CPLEX, while CASPO is able to retrieve the complete set of solutions.

**Table S3:** Case studies. 1) Nr. Edges: number of edges in the case-study; 2) Nr. Nodes: number of nodes; 3) Nr. Readouts: number of measurements; 4) Nr. Conditions: number of experimental conditions; 5) Time-points: time-points for each case-study (note that for cases with more than 2 time-points, the first 2 were considered for the analysis); 6) GA-50: time needed to run the analysis with GA with a population size of 50; 7) GA-100: time needed to run the analysis with GA with a population size of 100; 8) GA-300: time needed to run the analysis with GA with a population size of 300; 9) ASP: time needed for CASPO to enumerate all the solutions; 10) ILP: time needed for ILP to generate 1 optimal solution; 11) ILP-1000: time needed for ILP to enumerate a maximal number of 1000 optimal solutions.

| Case | Nr. Edges | Nr. Nodes | Nr. Readouts | Nr. Conditions | Time-points | GA-50 (seconds) | GA-100 (seconds) | GA-300 (seconds) | ASP (seconds) | ILP (seconds) | ILP-1000 (seconds) |
|---|---|---|---|---|---|---|---|---|---|---|---|
| **Toy-MSB 2009** | 10 | 8 | 4 | 5 | 0, 10 | 0.387 | 0.246 | 0.255 | 0.088 | 0.255 | 0.063 |
| **Toy-PCB** | 15 | 8 | 5 | 6 | 0, 30 | 2.100 | 6.532 | 12.107 | 0.015 | 0.038 | 0.068 |
| **Toy-PB** | 20 | 12 | 6 | 10 | 0, 2, 4, 6, …, 30 | 3.309 | 8.601 | 21.989 | 0.009 | 0.084 | 0.137 |
| **Liver-DREAM** | 62 | 17 | 7 | 25 | 0, 30 | 9.606 | 12.573 | 47.172 | 0.060 | 0.136 | 0.852 |
| **ExtLiver-PCB** | 99 | 31 | 15 | 64 | 0, 30 | 57.645 | 112.200 | 311.305 | 239.89 | 4.733 | 60.679 |
| **ExtLiver-PriHu** | 105 | 29 | 12 | 64 | 0, 30, 180 | 81.848 | 152.280 | 1123.622 | NA | 177.480 | 204.619 |
| **ExtLiver-BMC2012** | 108 | 31 | 15 | 64 | 0, 30 | 54.0378 | 127.433 | 725.962 | 273.66 | 33.933 | 127.827 |

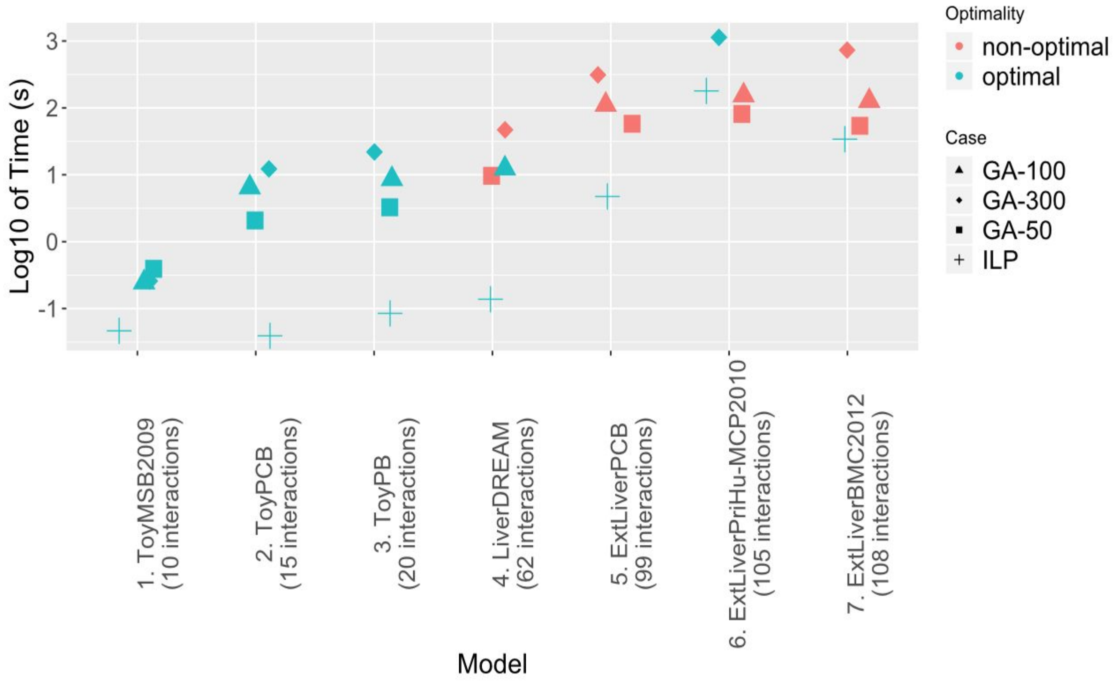

**Figure S1**. Computation time for different models ordered by the number of edges in the PKN. Cases are coloured red if the Integer Linear Programming (ILP) found a better optima than the Genetic Algorithm (GA). (GA-50: Genetic Algorithm with a population size of 50) ILP: Time needed for CPLEX to identify the first optimal solution).

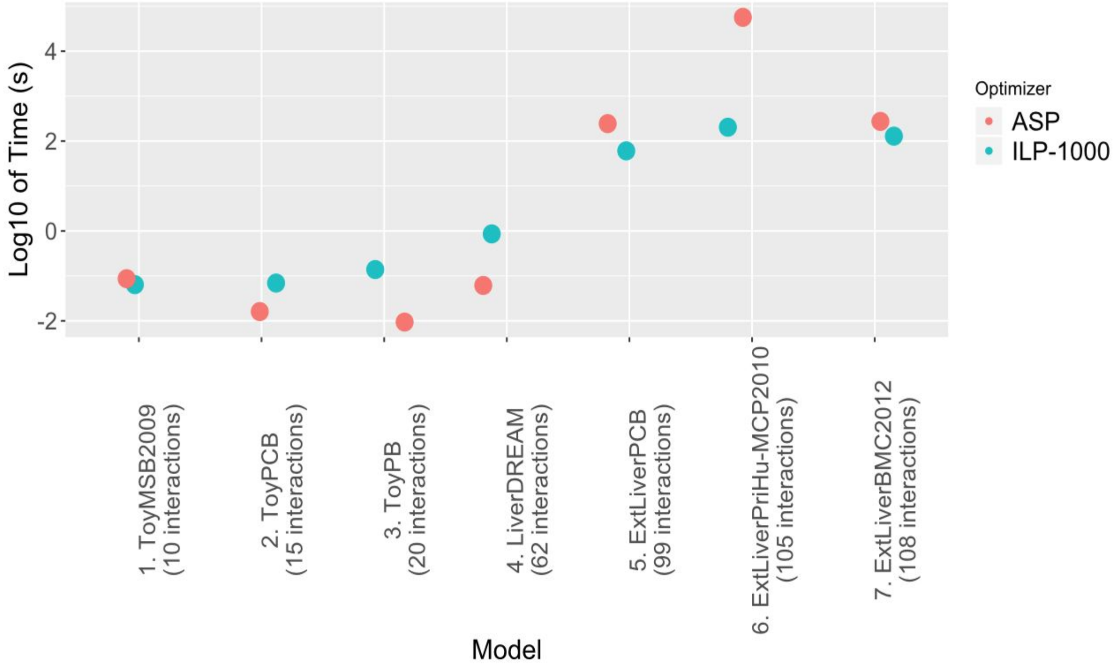

**Figure S2**. Computation time for different models ordered by the number of edges in the PKN. With red are shown the time needed for the Answer Set Programming (ASP) approach implemented in the Cell ASP Optimiser (CASPO) tool. CASPO generates all the optimal solutions. Green dots show the time needed for the ILP approach to enumerate a maximum of 1000 optimal solutions (ILP-1000). Note that for the case ExtLiverPriHu-MCP2010, CASPO was not able to enumerate the solutions even after 3 days running.

#### S1.4. Availability

The ILP implementation of CellNOpt is integrated within the main CellNOptR package. The documented R-package together with examples in the vignettes is available at https://github.com/saezlab/CellNOptR.

### Text S2. CellNOptR-MaBoSS

#### S2.1. Introduction

The Boolean logic models in CellNOpt are simulated with a deterministic simulator (Terfve et al. 2012). The new CellNOptR-MaBoSS package offers model structure optimisation based on population-based stochastic simulations. It integrates MaBoSS (Stoll et al. 2012), a stochastic simulator with CellNOptR (Terfve et al. 2012).

#### S2.2. CellNOpt-MaBoSS Pipeline

The CellNOpt-MaBoSS pipeline (from here on CNO-MaBoSS) builds on the CellNOptR package. It uses the same objective function (see S1.3.1 and equation (1)) and Genetic Algorithm to solve the optimisation problem of model training as CellNoptR. The major difference is at the model simulation level. CNO-MaBoSS simulates a population of networks asynchronously, then the trajectories are averaged in the population. This results in state trajectories that are continuous in [0, 1] and discrete in time.

This way CNO-MaBoSS simulations can handle time-course data and can capture oscillations. In this case, the oscillations can be seen as the result of cyclic attractors where the average node activity will finally be converged to a constant. When this happens, a stopping criteria has been implemented in the CNO-MaBoSS in order to avoid time-consuming simulations.

#### S2.3. Comparison of CNO-MaBoSS to CellNOptR, CNORode and FALCON

We compared CNO-MaBoSS to CellNOptR, FALCON and CNORode. The latter two methods aim to adjust continuous parameters on each species and interaction in the PKN associated with node responsiveness and strength of interactions respectively. CNORode implements logic-based Ordinary Differential Equations (ODE) within the CellNOptR framework as described in (Wittmann et al. 2009). FALCON (Landtsheer et al. 2017) on the other hand, assigns one parameter for each edge representing the flux of the signal from one species to the other.

To demonstrate the differences, we have applied the four methods to a published model by (MacNamara et al. 2012). It consists of 30 nodes and 33 edges (see example in the CNO-MaBoSS R-Package) and a negative feedback mechanism in the structure which accounts for the oscillatory behaviour of *NFkappaB*. This model is considered as a PKN in CellNOptR where non relevant nodes and edges can then be compressed (Saez-Rodriguez et al. 2009). Every experimental condition is a combination of stimulated inputs and inhibitors and measurements were considered to be obtained at time-points from 0 to 10 with intervals of length 2 between the timepoints.

For the first steady-state case-study only measurements at time-point 0 and 10 were considered and CNO-MaBoSS was compared to CellNOptR, FALCON and CNORode in terms of computational time and goodness of fit (Table S4.1). CNO-MaBoSS has a better fitting score than CellNOptR but it is slower. With CNORode, it is the opposite. CNORode has a better fitting score but is slower than the pipeline. FALCON performs well in both evaluations.

**Table S4.1.** Performances of CNO-MaBoSS and tested softwares (CellNOptR, CNORode and FALCON) - Case study 1. The score of the best solution of each tool is given by the MSE. Time execution of the tool to reveal the best solution is given in seconds (s).

| Software | MSE | Time (*s*) |
|---|---|---|
| **CNO-MaBoSS** | 0.0321 | 179.797 |
| **CellNOptR** | 0.0545 | 0.246 |
| **CNORode** | 0.0062 | 405.598 |
| **FALCON** | 0.0118 | 14.849 |

In the second case-study we made a comparison between CNO-MaBoSS and CNORode. In this case, the same model topology as before was considered, however, the measurements across all the time-courses were used for the analysis (Table S4.2). In terms of score, the results of the pipeline is worse than CNORode. However, the execution time of the pipeline is much faster than for CNORode optimisation while both are able to capture the observed oscillatory behaviour of the data due to the feedback mechanism in the network topology.

**Table S4.2.** Results and performance of CNO-MaBoSS compared to CNORode for the time course study (Case study 2).

| Software | MSE | Time (*s*) |
|---|---|---|
| **CNO-MaBoSS** | 0.0530 | 147.340 |
| **CNORode** | 0.0158 | 2354.000 |

The execution time of CNO-MaBoSS is much longer than the time needed to run CellNOptR and FALCON to obtain the best solution (optimised topology or parameter values) - several minutes against a few seconds. This is because CNO-MaBoSS uses stochastic simulations. However, CNO-MaBoSS can detect the feedback loops in a steady-state study and fit easily oscillations in a time-course study. The fitting quality of CNO-MaBoSS is worse compared to CNORode and FALCON, probably due to the fact that they use parameters to characterise the interactions while CNO-MaBoSS just identifies the presence or absence of these interactions. CNORode is even slower than the pipeline, due to the complexity of the equation in the framework, the number of parameters to fit and the combination of the global and local optimisation implemented in the enhanced Scatter Search method (eSSM) algorithm (Egea et al. 2009). However, CNORode gives better fitted models.

#### S2.3. Availability

CNO-MaBoSS has been implemented as an R-package and the tool together with examples can be found here: https://github.com/saezlab/CellNOptR-MaBoSS.

### Text S3. CNORprob

#### S3.1. CNORprob as an alternative to CNORfuzzy

To analyse logic models built upon quantitative data at a quasi steady-state, we previously applied the fuzzy logic formalism which allows to describe the changes in degree of membership (DOM) of node activity in the network within the range of 0 to 1 to capture semi-quantitative states (Morris et al. 2011). The computational pipeline to model and optimise fuzzy logic models was encoded in the CNORfuzzy package available at https://github.com/saezlab/cnorfuzzy.

Nevertheless, the CNORfuzzy framework requires the optimisation of 3 parameters per interaction which increases the computational time substantially. To account for this limitation, we applied a probabilistic logic network approach that can also represent the state of node in a semi-quantitative range between 0 to 1 while it contains only one probability parameter per interaction. Here, we compiled an R-package ‘CNORprob’ for the modelling and analysis of models in the probabilistic logic framework which was derived from the software suite ‘FALCON’ in Matlab (Landtsheer et al. 2017). FALCON was previously shown to perform the analyses faster than CNORfuzzy with a similar model fitting quality. The computational speeds of CNORprob and FALCON on the same case study are also highly comparable.

#### S3.2. Assignment of additive effects to logic gates

There is one major difference on the ground assumptions between the CellNOpt packages and CNORprob/FALCON that should be discussed. When there is more than one interaction coming to a node, the CellNOpt packages assume that these interactions will either take the OR or the AND gate. Alternatively, CNORprob considers the third case i.e. an additive effect without assigning the OR/AND gate in this scenario. This assignment gives more flexibility to the CNORprob framework as users could assign more types of reactions with a broader set of transfer functions.

#### S3.3. Availability

CNORprob is freely available at https://github.com/saezlab/CNORprob where users can find installation instructions and three real-case examples (Landtsheer et al. 2017; Trairatphisan et al. 2016; Lommel et al. 2016).

### Text S4. Dynamic-Feeder

#### S4.1. Introduction

CellNOpt derives models of signalling networks from a prior knowledge network (PKN). However, literature-constrained methods are limited to known interactions. Logic models are scalable models, but cannot cope with thousands of interactions reported in interaction databases, therefore the assessment of relevant prior knowledge always includes a selection of possible edges. This approach can be biased, incomplete or simply wrong.

Here we present Dynamic-Feeder to address the above issues. The Dynamic-Feeder represents a framework which identifies and incorporates new possible links to the PKN within a logic ordinary differential equation (ODE) formalism (Terfve et al. 2012; Wittmann et al. 2009). It is an extension of the previous CNORfeeder (Eduati et al. 2012) which was limited to the Boolean logic formalism.

The main difference between CNORfeeder and the Dynamic-Feeder is that the former finds possible interactions based on the perturbation data *before* fitting the model to data. Dynamic-Feeder assesses the quality of already trained models and sequentially integrates interactions to improve the fit, while using AIC and BIC criteria to avoid overfitting. Dynamic-Feeder can also search for new interactions from large signalling databases and weight the new integrated links based on the number of evidence from the literature.

#### S4.2. Dynamic-Feeder pipeline

The Dynamic-Feeder pipeline is shown in Figure S3.

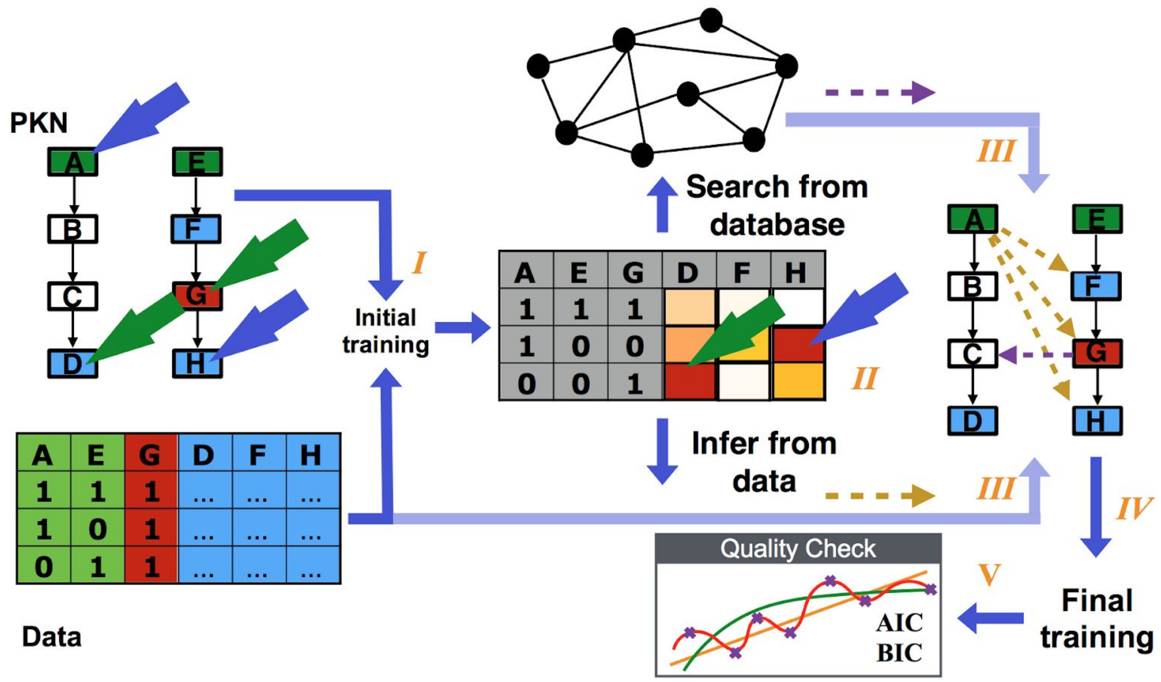

**Figure S3**. Dynamic-Feeder pipeline. *I*) Initial PKN is translated to dynamic models and trained to data. *II*) Identifying poorly fitted conditions. *III*) Add missing links to the PKN based on interaction databases (purple arrow) and links identified from the poorly fitted conditions (yellow arrows). *IV*) Integrating missing links to PKN and running final training to data results to better fits. *V*) Performing quality checks of the models to avoid overfitting through Akaike Information Criterion (AIC) and Bayesian Information Criterion (BIC) scores.

1. An initial dynamic analysis with CNORode is performed by training the PKN to data. This way we obtain the initial set of continuous parameters representing the dynamic features of nodes and edges in the PKN.

2. As a second step, we identify a set of measurements which are poorly fitted across experimental conditions. A poorly fitted measurement is defined as the one which has a mean square error (MSE) value worse than a threshold. The MSE is computed as the average of squares between the estimated measurement value by the model and the actual measured value. In this case, measurement $D$ is not fitted well for the experimental condition when $G$ is being inhibited, while $H$ is not predicted well by our model for the case when $A$ is being activated.

3. Poorly fitted nodes are an indication of a missing link in the PKN. After identifying the measurements that are poorly fitted, we can then identify possible interactions connecting the corresponding perturbed cues (activated and inhibited nodes) to these poorly fitted measurements. These new connections can be inferred from data alone by using the FEED algorithm (a data-driven interaction inference method described in (Eduati et al. 2012)) or searching from databases of interactions (within a certain path length limit), or both. As for the interaction resources, we particularly use OmniPath (Türei et al. 2016), a comprehensive collection of many publicly available curated resources about signalling pathways.

4. The final integrated model is then re-trained to the data where we can observe the effects of the new links over the fit of our new model predictions to the measurements.

5. Model selection methods can be applied to compare the quality of the original and the integrated network (i.e. Akaike Information Criterion (AIC) and Bayesian Information Criterion (BIC)).

There are some differences on how the FEED algorithm was applied in the Dynamic-Feeder compared to the original implementation which was strictly applied on a Boolean context. In the previous implementation, a link between a cue and any measurement was inferred if a stimulus or an inhibitor significantly affects an output protein. A protein was said to be significantly affected if its activity level changes by a quantity that exceeds the uncertainty of measurement on at least at one time-point. On the Dynamic-Feeder implementation however, we apply the FEED algorithm only on those nodes where the measurements are poorly fitted and the perturbed cues on the corresponding experimental conditions. This has an obvious computational advantage especially for the ODE case since we are integrating fewer links and only focusing on the possible ones which can improve the model predictions.

The goal of the training is to minimise the residual sum of squares of model predictions to the data with the minimum number of functional parameters. The objective function tries to balance the goodness of fit $\theta_f$ with the model size $\theta_s$ as described in (3) (Terfve et al. 2012):

$$min(\theta) = min(\theta_f + \theta_s) \qquad (3)$$

The goodness of fit is defined as MSE representing the deviations between measured data (m) and our model predictions (x):

$$\theta_f = \frac{1}{N} \sum_{i=1}^{n_s} \sum_{z=1}^{n_e} (m_{i,z} - x_{i,z})^2 \qquad (4)$$

For each species $i = \{1, 2, .., n_s\}$ and interaction $l = \{1, 2, .., n_r\}$ in the network the penalty of model size is formulated as:

$$\theta_s = \theta_s^\tau + \theta_s^k + \theta_s^n = \sum_{i=1}^{n_s} (\lambda_\tau^{PKN} + \lambda_\tau^{Database})\tau_i + \sum_{l=1}^{n_r} (\lambda_k^{PKN} + \lambda_k^{Database} + \lambda_k^{DDN})k_l + \sum_{l=1}^{n_r} (\lambda_n^{PKN} + \lambda_n^{Database} + \lambda_n^{DDN})n_l \qquad (5)$$

Here, the $\tau$ , $n$ and $k$ are the fitted ODE parameters associated with each dynamic state as described in (Krumsiek et al. 2010). These parameters can have some clear biological meaning. For example, $\tau_i$ parameter can be understood as the life-time of species $i$; while $k$ and $n$ are typically Hill parameters specific to each interaction $l$ in the network. While $k$ can be associated to strength of interaction (higher $k$ meaning stronger interaction and $k = 0$ meaning no link is present), the $n$ parameter can be associated with the cooperativity of the source and the target nodes in the interaction. In order to induce sparsity to the network, an L1-norm penalty term has been added to the objective function as shown in (5). Through the regularisation factors $\lambda_\tau$, $\lambda_k$ and $\lambda_n$, we can choose how to balance the goodness of fit with the size of the model. We can penalise the nodes and edges of the network and this way we also try to avoid over-fitting.Therefore, the higher penalty factor for the new members in the PKN will discourage the addition of new links, which do not provide a sensible contribution to the improvement of the overall fit.

Similar to CNORFeeder, we penalise the addition of the new links to the system by introducing another penalty factor to the objective function for each parameter associated with the new nodes and links. Normally, we should give higher priority to network components which are present in the original PKN than compared to the new components integrated via searching from signalling resources or inferred from data alone. Also regarding the newly integrated links, we can choose to give higher priority to the links inferred from the databases compared to the purely data-driven ones, since for the former links we have evidence from literature ($\lambda^{PKN} < \lambda^{Database} < \lambda^{DDN}$).

#### S4.3. Availability

Dynamic-Feeder has been integrated in the CNORfeeder (Eduati et al. 2012) package. The documented package together with examples can be found in https://github.com/saezlab/CellNOpt-Feeder. A real-case application of Dynamic-Feeder over the HPN-DREAM breast cancer dataset (Hill et al. 2016) has also been shown in (Gjerga et al. 2019).

### Text S5. Post-Hoc Analysis

#### S5.1. CNORprob

CNORprob implements post-optimisation analyses including systematic edge-knockout, systematic node-knockout, bootstrapping and local parameter sensitivity analyses (LPSA). Based on the results from post-hoc analyses, users could obtain the list of nodes and edges which are sensitive and insensitive to perturbations. Those which are insensitive to node and edge knockouts could potentially be removed from the model. In contrast, those which are sensitive, i.e. Akaike Information Criterion (AIC) value increases substantially after the edge/node removal, implies that the respective component of the model is vital for model fitting and could potentially be the key regulatory point. Bootstrapping analysis returns the information on the robustness of estimated weights against random noise while LPSA gives the clues whether the optimised parameter values are identifiable. All of these analyses offer an additional layer of information to understand and interpret the results from the optimised models in CNORprob.

#### S5.2. CNORode & CellNOptR

CNORode implements post-hoc analysis such as bootstrapping and cross-validations. Bootstrapping allows to estimate the robustness of the continuous parameters by performing the optimisation with random resampling (with replacement) of the experimental data multiple times. The feature is integrated within the CNORode package and it’s details are described in (Eduati et al. 2017).

On the other hand, in order to assess the predictive performance of our models and inspect how well these models can predict data outside of it's training set we have implemented cross-validation as another feature within the CNORode and CellNOptR framework. We have implemented three different re-sampling strategies about how we can split the training and the test set: either resampling of the data-points, experimental conditions or observable nodes.

#### S5.3. Availability

Examples of PostHoc analysis can be found on the corresponding documentations of CNORprob (https://github.com/saezlab/CNORprob), CNORode (https://github.com/saezlab/CNORode) and CellNOptR (https://github.com/saezlab/CellNOptR) packages.

### Text S6. ShinyCNOR

In order to facilitate the usage and exploration of functionality in the CellNOpt packages for new users, we also offer an interactive version of the three main CellNOpt packages as an R-Shiny application called ShinyCNOR. These include the CellNOpt, CNORprob and CNORode packages which are suitable for the optimisation of qualitative/discrete data at quasi steady-state, quantitative data at quasi steady-state and time-course quantitative data, respectively.

On the ShinyCNOR application, users can either upload their own data in the upload tab (network file in SIF format and data file in MIDAS format) or take a default example as provided by the software (Figure S4). Once the model and measurement files are uploaded or the example has been selected, the initial model structure and the measures are automatically visualised. In addition, the data table of the network and MIDAS files could also be displayed by clicking the corresponding tick-boxes.

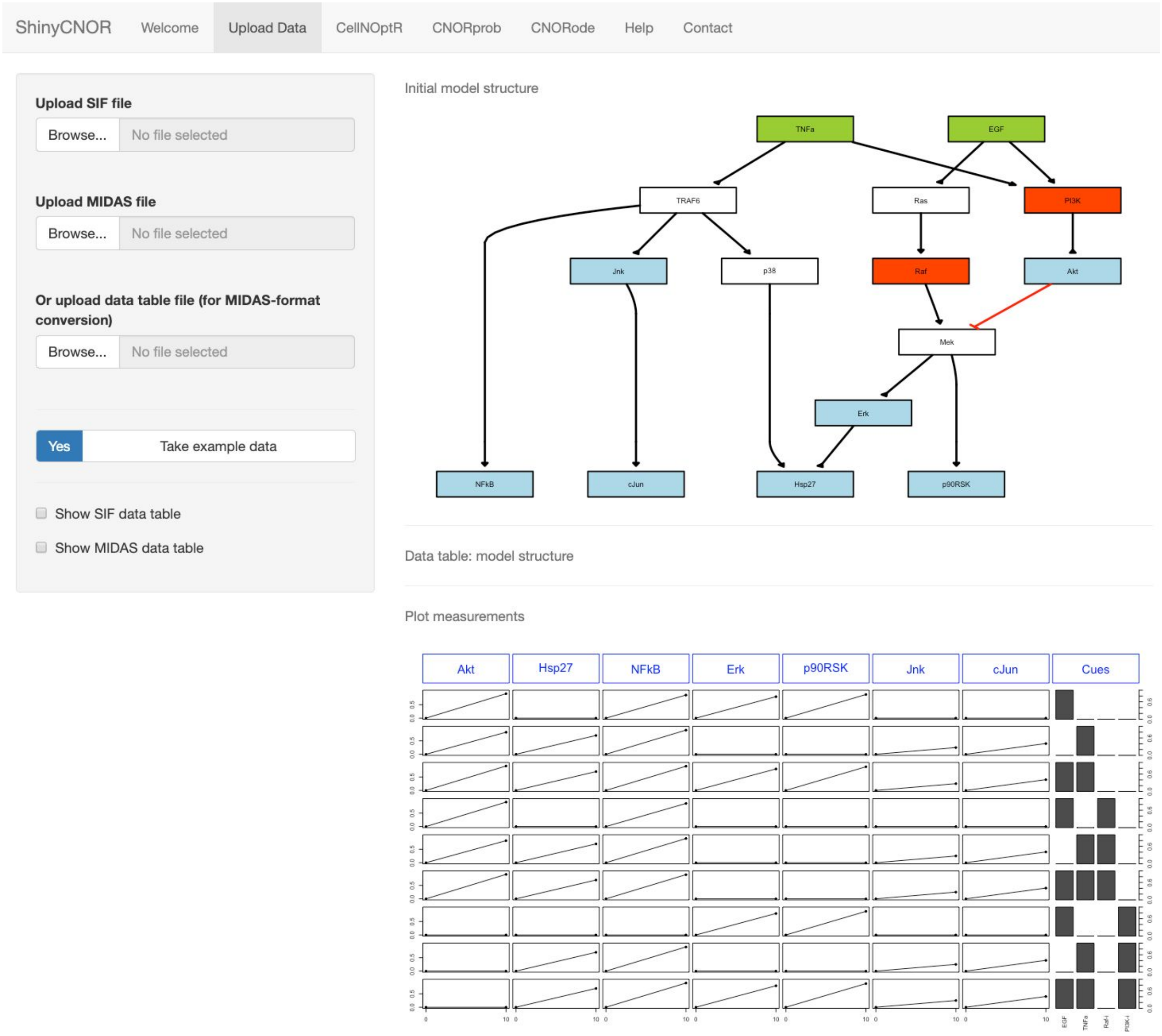

**Figure S4:** Graphical User Interface for uploading the model (from a SIF file) and data (from a MIDAS file).

In the CellNOpR tab, users can perform model compression and Boolean gates expansion as a pre-processing step (Saez-Rodriguez et al. 2009) on the initial network by selecting the provided options. After pre-processing, the network structure will be updated in real-time (Figure S5). Subsequently, users can click on the 'Optimise model' button to initiate the optimisation process. After the optimisation is done, the optimised network will be plotted and the model fitting quality will be displayed. Note that all optimisation parameters (by Genetic Algorithm) are provided on this tab so users can also customise these parameters.

In the CNORprob tab (Figure S6), users can perform a pre-processing of the initial network similar to the CellNOptR tab. Nevertheless, given that the consideration of Boolean gate assignment in CNORprob is different compared to the other CellNOpt packages (see Supplementary Text 3 - CNORprob), the assignment of OR-gate expansion will replace the additive effect of edge combination which is assigned by default. Then, users can click the

‘Run CNORprob’ button to execute the optimisation. After the optimisation, the optimised network will be plotted together with the edge weight/probability as well as the model fitting quality in another figure. In addition, users can also run post-hoc analyses including edge/node knock-out, (local) parameter sensitivity analysis and bootstrapping analysis by ticking the corresponding options and click on the “Run Post-hoc analysis” button. The results from the analysis will then be saved in the ‘Results’ folder on the current working directory in the R-session.

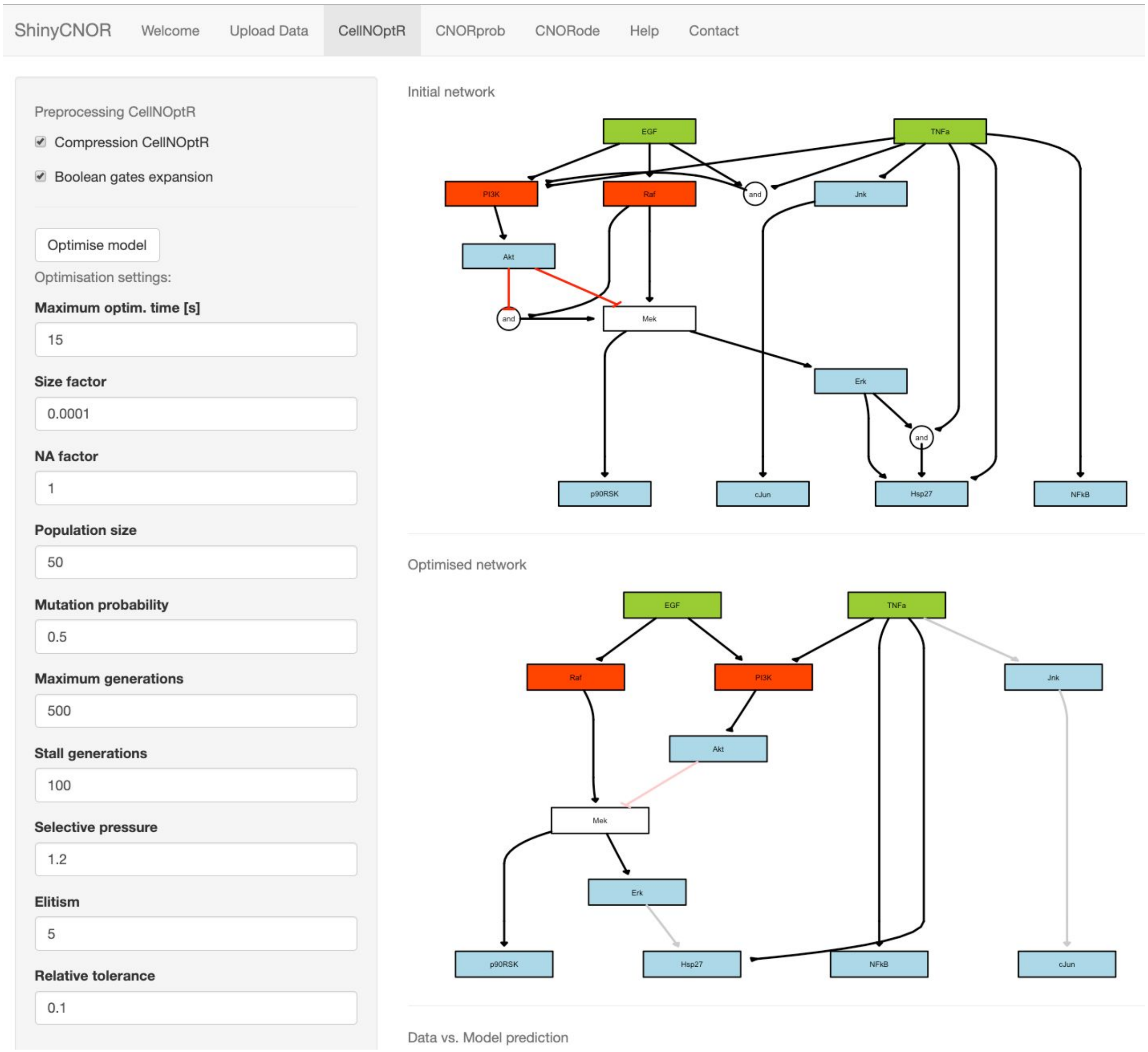

**Figure S5:** Graphical User Interface for the pre-processing of the model (compression and expansion).

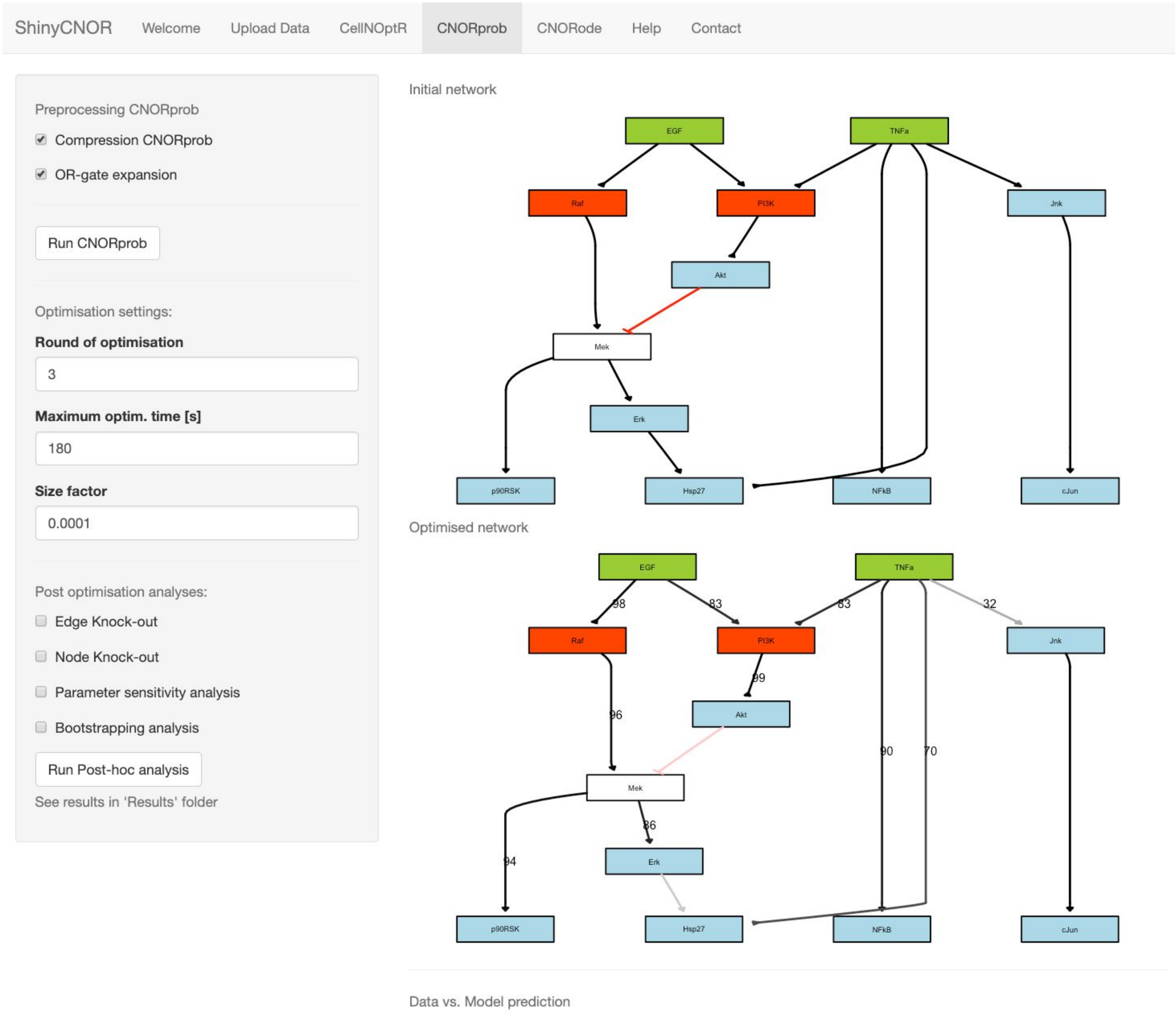

**Figure S6:** Graphical User Interface for analysis with CNORprob.

In the CNORode tab (Figure S7), users can perform the compression of the initial network but not Boolean gate expansion which does not fit to the Logic-ODE framework (either the OR gate or the AND gate has to be assigned for each interaction beforehand to prevent ambiguity during the optimisation). Also three different types of transfer function can be selected i.e. Hill (transfer function 2), Normalised Hill (transfer function 3) (Wittmann et al. 2009) and Inverse normalised Hill (transfer function 4) (Eduati et al. 2017) which might be suited for different types of optimisation problem. The bound of parameters in the Logic-ODE framework i.e. $k$, $n$ and $\tau$ can also be directly scaled within the tool together with the termination criteria of the optimisation. By running the optimisation by clicking the button 'Run CNORode', the optimised model with fitting quality figures are displayed. Furthermore, users can also download the script to run the optimisation on a local R-session by clicking the 'Download script' button in case the optimisation problem is very large and could not be handled well within the Shiny application.

For additional information on the usage and trouble-shooting of each tab, users can consult the ‘Help’ tab. All issues can be reported to our GitHub issue page.

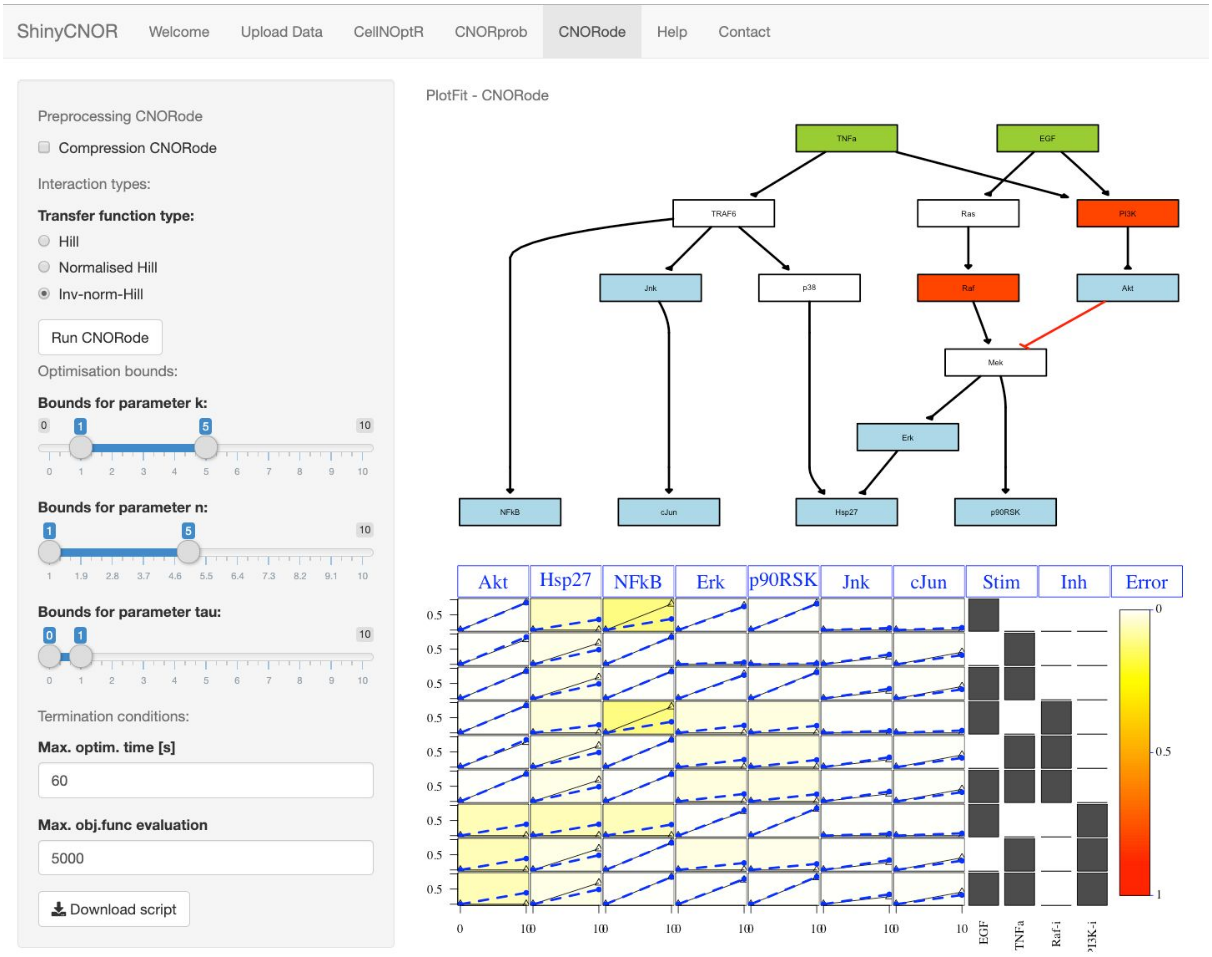

**Figure S7:** Graphical User Interface for analysis with CNORode.

#### S6.1. Availability

The software can be run locally after the installation from GitHub (https://github.com/saezlab/ShinyCNOR) or online through the R-Shiny application (https://saezlab.shinyapps.io/shinycnor/).

### References:

Albert, István, Juilee Thakar, Song Li, Ranran Zhang, and Réka Albert. 2008. “Boolean Network Simulations for Life Scientists.” *Source Code for Biology and Medicine* 3 (November): 16.

Alexopoulos, Leonidas G., Julio Saez-Rodriguez, Benjamin D. Cosgrove, Douglas A.

Lauffenburger, and Peter K. Sorger. 2010. “Networks Inferred from Biochemical Data Reveal Profound Differences in Toll-like Receptor and Inflammatory Signaling between Normal and Transformed Hepatocytes.” *Molecular & Cellular Proteomics: MCP* 9 (9): 1849–65.

Batt, Grégory, Bruno Besson, Pierre-Emmanuel Ciron, Hidde de Jong, Estelle Dumas, Johannes Geiselmann, Regis Monte, et al. 2012. “Genetic Network Analyzer: A Tool for the Qualitative Modeling and Simulation of Bacterial Regulatory Networks.” *Methods in Molecular Biology* 804: 439–62.

Benque, David, Sam Bourton, Caitlin Cockerton, Byron Cook, Jasmin Fisher, Samin Ishtiaq, Nir Piterman, Alex Taylor, and Moshe Y. Vardi. 2012. “Bma: Visual Tool for Modeling and Analyzing Biological Networks.” *Computer Aided Verification*. https://doi.org/10.1007/978-3-642-31424-7_50.

Chaouiya, Claudine, Duncan Bérenguier, Sarah M. Keating, Aurélien Naldi, Martijn P. van Iersel, Nicolas Rodriguez, Andreas Dräger, et al. 2013. “SBML Qualitative Models: A Model Representation Format and Infrastructure to Foster Interactions between Qualitative Modelling Formalisms and Tools.” *BMC Systems Biology* 7 (December): 135.

Chaouiya, Claudine, Aurélien Naldi, and Denis Thieffry. 2012. “Logical Modelling of Gene Regulatory Networks with GINsim.” *Methods in Molecular Biology* 804: 463–79.

Chun, Hyonho, Jia Kang, Xianghua Zhang, Minghua Deng, Haisu Ma, and Hongyu Zhao. 2011. “Reverse Engineering of Gene Regulation Networks with an Application to the DREAM4 in Silico Network Challenge.” *Handbook of Statistical Bioinformatics*. https://doi.org/10.1007/978-3-642-16345-6_22.

Dorier, Julien, Isaac Crespo, Anne Niknejad, Robin Liechti, Martin Ebeling, and Ioannis Xenarios. 2016. “Boolean Regulatory Network Reconstruction Using Literature Based Knowledge with a Genetic Algorithm Optimization Method.” *BMC Bioinformatics* 17 (1): 410.

Di Cara, A., Garg, A., De Micheli, G. *et al.* Dynamic simulation of regulatory networks using SQUAD. *BMC Bioinformatics* 8, 462 (2007). https://doi.org/10.1186/1471-2105-8-462

Eduati, Federica, Javier De Las Rivas, Barbara Di Camillo, Gianna Toffolo, and Julio Saez-Rodriguez. 2012. “Integrating Literature-Constrained and Data-Driven Inference of

Signalling Networks.” *Bioinformatics* 28 (18): 2311–17.

Eduati, Federica, Victoria Doldàn-Martelli, Bertram Klinger, Thomas Cokelaer, Anja Sieber, Fiona Kogera, Mathurin Dorel, Mathew J. Garnett, Nils Blüthgen, and Julio Saez-Rodriguez. 2017. “Drug Resistance Mechanisms in Colorectal Cancer Dissected with Cell Type–Specific Dynamic Logic Models.” *Cancer Research*. https://doi.org/10.1158/0008-5472.can-17-0078.

Eduati, Federica, Patricia Jaaks, Jessica Wappler, Thorsten Cramer, Christoph A. Merten, Mathew J. Garnett, and Julio Saez-Rodriguez. 2020. “Patient-specific Logic Models of Signaling Pathways from Screenings on Cancer Biopsies to Prioritize Personalized Combination Therapies.” *Molecular Systems Biology*. https://doi.org/10.15252/msb.20188664.

Egea, Jose A., Eva Balsa-Canto, María-Sonia G. García, and Julio R. Banga. 2009. “Dynamic Optimization of Nonlinear Processes with an Enhanced Scatter Search Method.” *Industrial & Engineering Chemistry Research*. https://doi.org/10.1021/ie801717t.

Gjerga, Enio, Panuwat Trairatphisan, Attila Gabor, and Julio Saez-Rodriguez. 2019. “Literature and Data-Driven Based Inference of Signalling Interactions Using Time-Course Data.” *IFAC-PapersOnLine*. https://doi.org/10.1016/j.ifacol.2019.12.235.

Guziolowski, Carito, Santiago Videla, Federica Eduati, Sven Thiele, Thomas Cokelaer, Anne Siegel, and Julio Saez-Rodriguez. 2013. “Exhaustively Characterizing Feasible Logic Models of a Signaling Network Using Answer Set Programming.” *Bioinformatics* 29 (18): 2320–26.

Helikar, Tomáš, Bryan Kowal, Sean McClenathan, Mitchell Bruckner, Thaine Rowley, Alex Madrahimov, Ben Wicks, Manish Shrestha, Kahani Limbu, and Jim A. Rogers. 2012. “The Cell Collective: Toward an Open and Collaborative Approach to Systems Biology.” *BMC Systems Biology* 6 (August): 96.

Hill, Steven M., Laura M. Heiser, Thomas Cokelaer, Michael Unger, Nicole K. Nesser, Daniel E. Carlin, Yang Zhang, et al. 2016. “Inferring Causal Molecular Networks: Empirical Assessment through a Community-Based Effort.” *Nature Methods* 13 (4): 310–18.

Krumsiek, Jan, Sebastian Pölsterl, Dominik M. Wittmann, and Fabian J. Theis. 2010.

“Odefy--from Discrete to Continuous Models.” *BMC Bioinformatics* 11 (May): 233.

Landtsheer, Sébastien De, Sébastien De Landtsheer, Panuwat Trairatphisan, Philippe Lucarelli, and Thomas Sauter. 2017. “FALCON: A Toolbox for the Fast Contextualization of Logical Networks.” *Bioinformatics*. https://doi.org/10.1093/bioinformatics/btx380.

Lommel, Maiti J., Panuwat Trairatphisan, Karoline Gäbler, Christina Laurini, Arnaud Muller, Tony Kaoma, Laurent Vallar, Thomas Sauter, and Elisabeth Schaffner-Reckinger. 2016. “L-Plastin Ser5 Phosphorylation in Breast Cancer Cells and in Vitro Is Mediated by RSK Downstream of the ERK/MAPK Pathway.” *FASEB Journal: Official Publication of the Federation of American Societies for Experimental Biology* 30 (3): 1218–33.

MacNamara, Aidan, Camille Terfve, David Henriques, Beatriz Peñalver Bernabé, and Julio Saez-Rodriguez. 2012. “State-Time Spectrum of Signal Transduction Logic Models.” *Physical Biology* 9 (4): 045003.

Mitsos, Alexander, Ioannis N. Melas, Paraskeuas Siminelakis, Aikaterini D. Chairakaki, Julio Saez-Rodriguez, and Leonidas G. Alexopoulos. 2009. “Identifying Drug Effects via Pathway Alterations Using an Integer Linear Programming Optimization Formulation on Phosphoproteomic Data.” *PLoS Computational Biology* 5 (12): e1000591.

Morris, Melody K., Julio Saez-Rodriguez, David C. Clarke, Peter K. Sorger, and Douglas A. Lauffenburger. 2011. “Training Signaling Pathway Maps to Biochemical Data with Constrained Fuzzy Logic: Quantitative Analysis of Liver Cell Responses to Inflammatory Stimuli.” *PLoS Computational Biology* 7 (3): e1001099.

Müssel, Christoph, Martin Hopfensitz, and Hans A. Kestler. 2010. “BoolNet--an R Package for Generation, Reconstruction and Analysis of Boolean Networks.” *Bioinformatics* 26 (10): 1378–80.

Naldi, Aurélien, Céline Hernandez, Wassim Abou-Jaoudé, Pedro T. Monteiro, Claudine Chaouiya, and Denis Thieffry. 2018. “Logical Modeling and Analysis of Cellular Regulatory Networks With GINsim 3.0.” *Frontiers in Physiology* 9 (June): 646.

Rodriguez, Ana, Isaac Crespo, Ganna Androsova, and Antonio del Sol. 2015. “Discrete Logic Modelling Optimization to Contextualize Prior Knowledge Networks Using PRUNET.” *PloS One* 10 (6): e0127216.

Saez-Rodriguez, Julio, Leonidas G. Alexopoulos, Jonathan Epperlein, Regina Samaga, Douglas A. Lauffenburger, Steffen Klamt, and Peter K. Sorger. 2009. “Discrete Logic Modelling as a Means to Link Protein Signalling Networks with Functional Analysis of Mammalian Signal Transduction.” *Molecular Systems Biology*. https://doi.org/10.1038/msb.2009.87.

Schwab, Julian D., and Hans A. Kestler. 2018. “Automatic Screening for Perturbations in Boolean Networks.” *Frontiers in Physiology* 9 (April): 431.

Stoll, Gautier, Barthélémy Caron, Eric Viara, Aurélien Dugourd, Andrei Zinovyev, Aurélien Naldi, Guido Kroemer, Emmanuel Barillot, and Laurence Calzone. 2017. “MaBoSS 2.0: An Environment for Stochastic Boolean Modeling.” *Bioinformatics* 33 (14): 2226–28.

Terfve, Camille, Thomas Cokelaer, David Henriques, Aidan MacNamara, Emanuel Goncalves, Melody K. Morris, Martijn van Iersel, Douglas A. Lauffenburger, and Julio Saez-Rodriguez. 2012. “CellNOptR: A Flexible Toolkit to Train Protein Signaling Networks to Data Using Multiple Logic Formalisms.” *BMC Systems Biology* 6 (October): 133.

Tognetti, Marco, Attila Gabor, Mi Yang, Valentina Cappelletti, Jonas Windhager, Konstantina Charmpi, Natalie de Souza, Andreas Beyer, Julio Saez-Rodriguez, Bernd Bondemiller. “Deciphering the Signaling Network Landscape of Breast Cancer Improves Drug Sensitivity Prediction.” https://doi.org/10.1101/2020.01.21.907691.

Trairatphisan, Panuwat, Andrzej Mizera, Jun Pang, Alexandru Adrian Tantar, and Thomas Sauter. 2014. “optPBN: An Optimisation Toolbox for Probabilistic Boolean Networks.” *PloS One* 9 (7): e98001.

Trairatphisan, Panuwat, Monique Wiesinger, Christelle Bahlawane, Serge Haan, and Thomas Sauter. 2016. “A Probabilistic Boolean Network Approach for the Analysis of Cancer-Specific Signalling: A Case Study of Deregulated PDGF Signalling in GIST.” *PloS One* 11 (5): e0156223.

Traynard, Pauline, Luis Tobalina, Federica Eduati, Laurence Calzone, and Julio Saez-Rodriguez. 2017. “Logic Modeling in Quantitative Systems Pharmacology.” *CPT: Pharmacometrics & Systems Pharmacology*. https://doi.org/10.1002/psp4.12225.

Türei, Dénes, Tamás Korcsmáros, and Julio Saez-Rodriguez. 2016. “OmniPath: Guidelines and Gateway for Literature-Curated Signaling Pathway Resources.” *Nature Methods* 13

(12): 966–67.

“Website.” n.d. Accessed February 21, 2020. Di Cara, A., Garg, A., De Micheli, G. et al. Dynamic simulation of regulatory networks using SQUAD. BMC Bioinformatics 8, 462 (2007). https://doi.org/10.1186/1471-2105-8-462.

Wittmann, Dominik M., Jan Krumsiek, Julio Saez-Rodriguez, Douglas A. Lauffenburger, Steffen Klamt, and Fabian J. Theis. 2009. “Transforming Boolean Models to Continuous Models: Methodology and Application to T-Cell Receptor Signaling.” *BMC Systems Biology* 3 (September): 98.
